# Accurate evaluation of neural sensitivity to weak alternating electric fields requires protocols accounting for input dependence

**DOI:** 10.1101/2025.11.24.690108

**Authors:** Gabriel Gaugain, Julien Modolo, Denys Nikolayev

## Abstract

*Background:* Weak alternating electric fields (∼1 V/m) modulate nervous system activity. Yet, the exact mechanism by which such low amplitude electric fields can modulate neural activity is still unknown despite the use of specific protocols aiming at quantifying neural sensitivity. *Objective:* To characterize how the measurement protocol can impact neural sensitivity to weak electric fields and bias *in vivo* sensitivity. *Methods:* We considered a variety of somatic clamp stimulation to drive the activity of biophysical morpho-realistic reconstructed neurons during extracellular alternating stimulation to quantify the sensitivity to the field depending on the nature of cells’ activity. *Results:* Cells sensitivity to alternating current stimulation depended on the type of input used to drive their activity, with a different frequency response for each protocol, with a trend for inhibitory neurons to be more sensitive to higher stimulation frequencies. Even with the same clamp protocol, sensitivity depended on the statistics of the input used. *Significance:* Neuronal sensitivity to alternating current stimulation is highly input-dependent, which has been largely neglected so far, and depends on the current statistic of the received inputs, and is not reliably represented by simple current clamps protocols.

## Introduction

Modulating brain activity to achieve a desirable effect on behavior or to relieve the symptoms associated with neurological disorders is key in neuroscience. Alternating electric fields are candidates to influence brain oscillations, being delivered non-invasively by modalities such as transcranial alternating current or magnetic stimulation (tACS/TAMS), or invasively through electro-cortical stimulation. While many studies performed *in-vitro* measurements (Deans et al., 2007; Reato et al., 2010; Anastassiou et al., 2011a) and *in-vivo* measurements (Fröhlich & McCormick, 2010; Johnson et al., 2020; Krause et al., 2019, 2022a; Ozen et al., 2010), knowing precisely and *a priori* the impact of alternating stimulation on the characteristics on neuronal activity remains a challenge.

While previous studies have pointed to alternating currents (AC) – and the related alternating electric fields (AEF) – effects being related to network effects (Francis et al., 2003; Zhao et al., 2024), precisely quantifying their impact at the single cell level is crucial for further theoretical network developments. Neocortical columns are indeed made of thousands of neurons with diverse types, and simulating all of these neurons is almost computationally intractable since it requires considerable computing time and resources. This technological achievement was conducted by a collaborative effort (Markram et al., 2015; Isbister et al., 2024), but remains on the fringes of typical studies. Furthermore, studying AC effects requires testing a large parameter space (amplitude, frequency, and possibly duration of stimulation). The translation of AC effects at the detailed single-cell level to simplified models, which could enable the testing of further network effects.

Recent computational studies highlighted a similar threshold for AC significant phase entrainment at the single cell level (Lee et al., 2024; Tran et al., 2022a), contrasting with the lower effects in isolated cells reported in (Francis et al., 2003). In (Lee et al., 2024) the model is directly based on measurements while the one developed in (Aberra et al., 2018) and used in (Tran et al., 2022a) is adapted from the Blue Brain Project reconstructed cell database (Ramaswamy et al., 2015) to incorporate myelin, therefore changing the mechanisms adapted from measurements. The original cells were used in (Gaugain et al., 2025), but did not include the axonal tree, commonly modeled as a stub similar to (Lee et al., 2024). Not only did the cell models change across these studies, but the stimulus to drive their activities changed subsequently with a monophasic current clamp (Lee et al., 2024), one 50 Hz-synapse on the apical tree (Tran et al., 2022a), and the full set of reconstructed synapses (Gaugain et al., 2025). The difference in the nature of activity could further change their sensitivity to AEF.

Here, we investigated the impact of different protocols to drive detailed single-cells’ activity on their sensibility to AC. The models reported by the Blue Brain Project (Markram et al., 2015), and previously used (Gaugain et al., 2025), with current clamps as well as dynamic clamps to generate supra-threshold activities during exposition to AEF to quantify cells sensitivity depending on the nature of the activity/inputs.

## Materials and Methods

### Neuron models

Morphologically reconstructed neurons published in (Markram et al., 2015) were used as realistic models of neuronal geometry. These models were originally reconstructed from images of juvenile rat somatosensory cortex neurons and segmented into compartments compatible with the NEURON simulator (Hines & Carnevale, 1997). 13 Hodgkin–Huxley type channels were optimized to fit the data from electrophysiological recordings. In-house Python scripts were developed to control all simulations and are available on Github. Here, we selected five cell models considering a layer 5 thick tufted with early bifurcation pyramidal cell (PC) and four layer 4 inhibitory cells among vaso-intestinal peptide (VIP), somatostatin (SST) and parvalbumin (PV) expressing interneurons, namely one bipolar cell, one Martinotti cell, and one large basket cell.

### Neural activity source

Neural activity was driven either by current injection through a current clamp or a dynamic clamp, which is not novel but rarely used. This method requires a real-time computing unit and operating system as well as two clamps. Therefore, the current clamp technique is more often used. To test if the nature of neural activity impacts its sensitivity to weak electric fields, we tested four different cases. First, we considered neural activity driven by a constant monophasic current (Anastassiou et al., 2011b; Lee et al., 2024).

Second, an additional white noise was added to the constant current amplitude, with the related standard deviation 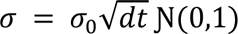, where Ɲ(0,1) denotes the normal distribution, *dt* is the time step of NEURON simulations, and *σ*_0_ the scaling factor.

Third, we considered a current injection with an amplitude driven by an artificial excitatory synapse consisting in a biexponential synapse of 2 ms rise time constant and 10 ms decaying time constant triggered by a random Poisson’s process. The firing rate of this process was set to 10, 20, 50, 100, and 3000 Hz to test the impact on stimulus variation frequency while the weight, or post-synaptic potential amplitude, was used to control the firing rate of the cell. This case corresponds to the one reported *in silico* in (Tran et al., 2022a), and was also previously used in (Chance et al., 2002) with higher frequencies to study gain modulation through synaptic input.

Finally, a synaptic barrage protocol which would involve a dynamic clamp setup (Sharp et al., 1992, 1993), such as the one described in (Destexhe et al., 2001; Destexhe & Paré, 1999) was used to trigger a desired activity while getting closer to the *in vivo* conditions of the cells. These synaptic barrage stimuli consisted of combined excitatory and inhibitory synapses with associated conductances following a stochastic random process similar to the Ornstein-Uhlenbeck process (Destexhe et al., 2001), described as follow:

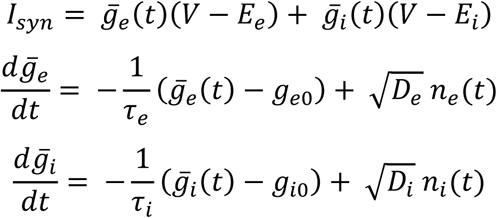

Where *g̅*_*e*_(*t*) and *g̅*_*i*_ (*t*) are excitatory and inhibitory dynamic conductance with their related reversing potential *E*_*e*_ and *E*_*i*_, and average conductance *g*_*e*0_ and *g*_*i*0_, *τ*_*e*_ and *τ*_*i*_ their time constant, *D*_*e*_and *D*_*i*_are noise ‘diffusion’, and *n*_*e*_(*t*)/*n*_*i*_(*t*) are normally distributed gaussian noise. To reproduce *in vivo* like synaptic bombardment, excitatory and inhibitory conductances shared the same mean and standard deviations for inhibitory cells while inhibitory conductances were 4 times larger than excitatory ones for the pyramidal cell accordingly (Salkoff et al., 2015; Shu et al., 2003).

For all protocols, the stimulation was set such that the cells exhibited a 10 Hz firing rate, often associated with alpha rhythm and enabled comparison with (Tran 2022, Gaugain 2025). All the parameters are presented in Table S1.

### Alternating electric field stimulation

Extracellular alternating electric field stimulation was modeled using the *extracellular* mechanism included in the NEURON simulator (Hines & Carnevale, 1997) together with the *xtra* mechanism to control stimulation waveform. The quasi-uniform approximation (Bikson et al., 2013) was used, and the electric field was set to be uniform and oriented in the *y*-direction, which was the somato-dendritic axis of pyramidal cells. The resulting extracellular potential was calculated as *e*_*i*_^*extra*^(*t*) = *E*(*t*) ⋅ *x*_*i*_, where *e*_*i*_^*extra*^(*t*) is the extracellular potential at the *i*^*th*^ compartment at time *t*, *x*_*i*_ its coordinates, *E*(*t*) = *E*_0_*u*_*y*_ ⋅ *sin*(2*πft*) the electric field vector with *E*_0_ the electric field amplitude in V/m, and *f* its frequency.

### Phase entrainment quantification

Spike events were extracted using the *eFEL Library* (Geit et al., 2018), and neural entrainment due to alternating electric field was assessed by the phase locking value (PLV) (Vinck et al., 2010) defined as:

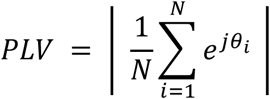

where N is the number of spike events, and *θ*_*i*_ the tACS waveform phase at which *i*^*th*^ spike occurs. This metric quantifies spike synchrony relative to the alternating stimulus with a value ranging from 0 (no synchronization) to 1 (perfect synchronization). The use of PLV is common in neural entrainment assessment (Johnson et al., 2020; Krause et al., 2019; Tran et al., 2022b) (Krause et al., 2019; Johnson et al., 2020; Tran et al., 2022a) while being a biased estimate (Vinck et al., 2010), since it is tightly linked to the commonly performed Rayleigh test that was used to test significant deviation from uniform phase distribution.

## Results

### Pyramidal cells exhibit input-dependent sensitivity to electric fields

L5 PC showed a clear input-dependent sensitivity to weak alternating electric field (wAEF) as depicted in Figure 1, where results for control and 10 V/m stimulation are depicted for comparison. Using monophasic stimulus to drive the selected L5 PC activity made the cell oversensitive to wAEF, with all spikes occurring at the same electric field phase (figure 1D, left panel). This could be due to the already regular activity of the cell generated by monophasic current clamps, which is easier to bias/entrain with an external stimulation. This was also observed for the gaussian protocol with a weaker effect, as the additional noise softly weakened the regular spiking of the cell. However, the two dynamic clamp protocols with a single synapse and synaptic bombardment generated a much more complex and irregular spiking pattern due to the random occurrence of synaptic events. The activity generated by a 50 Hz single synapse was significantly entrained by a 10 V/m 10-Hz AEF while the control activity was not phase locked at 10 Hz. Synaptic bombardment generated activity was not entrained even with a 10 V/m stimulation, showing that even a high level of EF could fail to entrain activity depending on the nature/statistics of cells’ activity.

**Figure 1.**
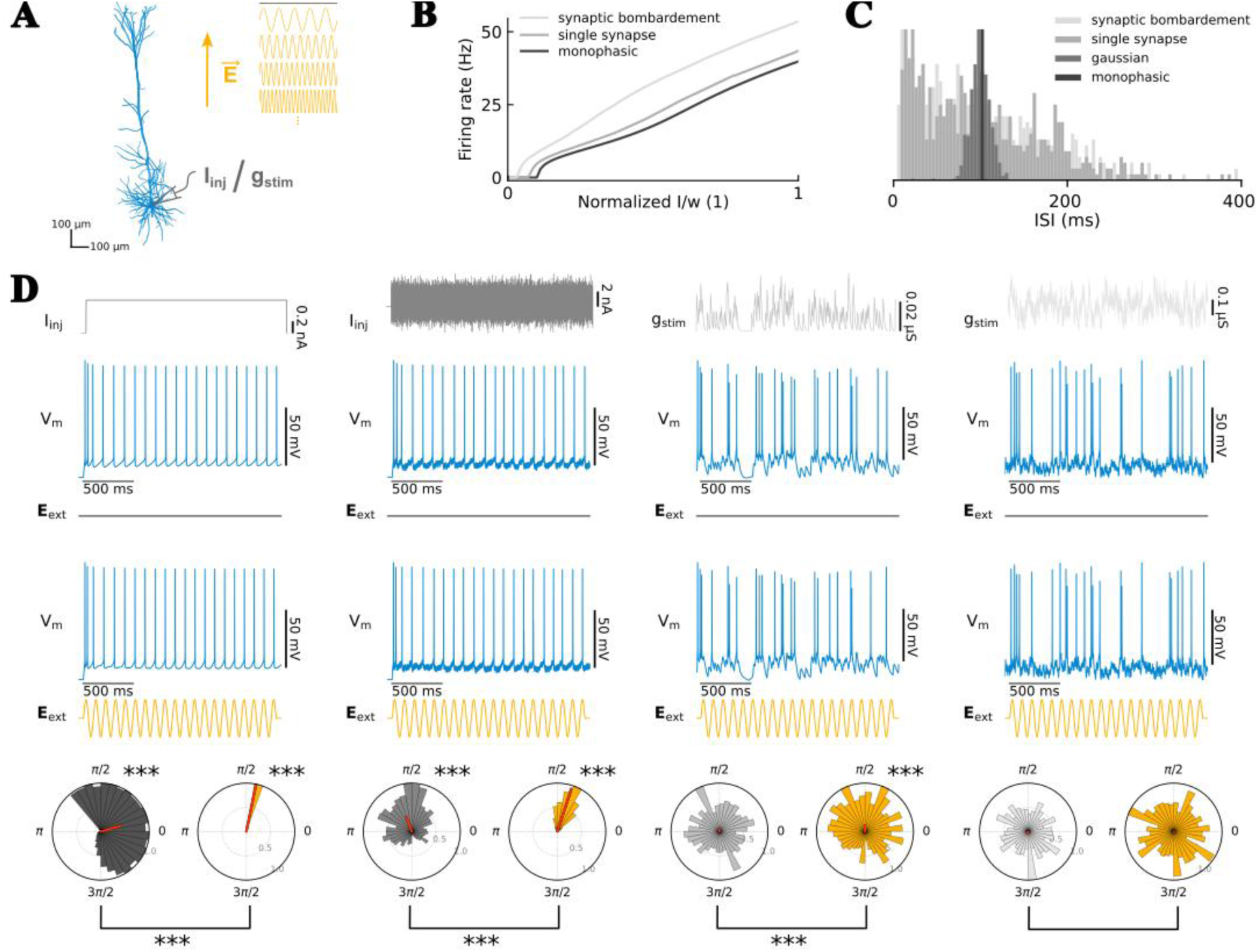
Layer 5 pyramidal cell response to 10-Hz tACS in the four different clamp stimulation protocols. A: Pyramidal cell’s morphology with clamp and electric field stimulation illustrated. B: Firing rate curves for the different protocols (monophasic ∼ gaussian noise) and the respective induced inter-spike intervals distributions. C: Traces of the four different clamp protocols without alternating electric field (top traces) and during a 10V/m electric field stimulation (bottom traces). D: Associated polar plots (grey shades for control and blue ones for the electric field stimulation condition), with Rayleigh test significances denoted in their associated right corner (***: p<0.001) and mean vector in red. A non-parametric permutation test was performed for each couple to test if tACS significantly increased PLV (***: p<0.001).

Furthermore, the investigation of frequency dependence of cell’s activity entrainment to AC demonstrated the ability to entrain cell’s activity at different stimulation frequencies for both monophasic and monophasic + gaussian noise current clamp protocols (Figure 2A & B). Generated activities were significantly entrained at frequencies multiple of the intrinsic spiking frequency – here, 10 Hz – showing an Arnold tongue-like response. Entrainment decreased faster as the multiple increased for the extra gaussian noise protocol, and lower amplitude of the monophasic current. Interestingly, an exception was observed at 1 V/m for gaussian noise stimulus for which the entrainment to the AES was found higher at the second harmonic (20 Hz). These tendencies disappeared for both dynamic clamp protocols, in which no clear relationships between stimulation frequency and entrainment were observed (Figure 2C & D). Therefore, this Arnold tongue-like response to AC depends critically on the protocol used to drive activity.

**Figure 2.**
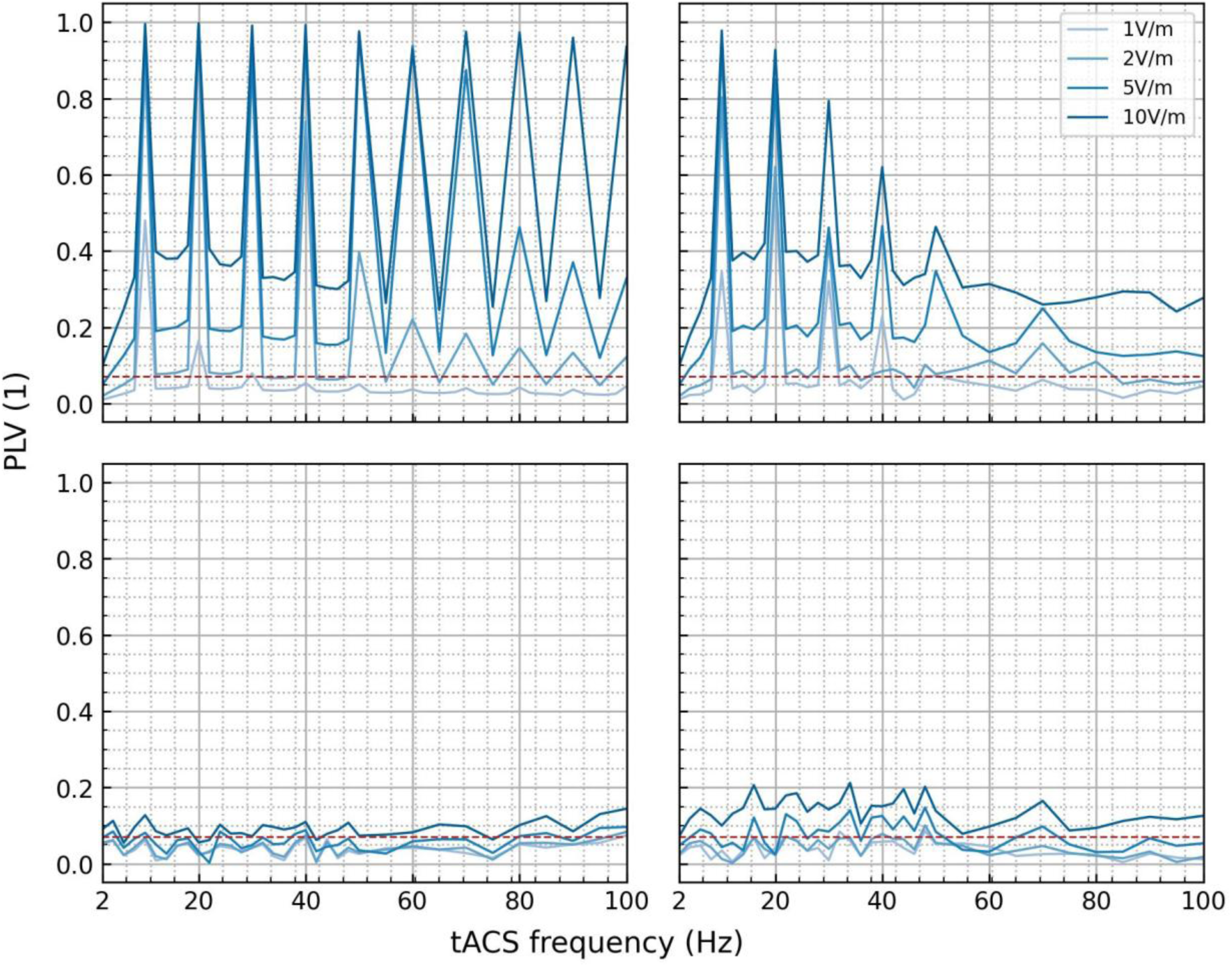
Layer 5 pyramidal cell’s response to the four tested clamp protocols during alternating electric field stimulation at 1, 2, 5 and 10 V/m with various stimulation frequencies. Phase locking values quantified entrainment (y-axis). Monophasic (top left) and monophasic + gaussian noise (top right) currents clamp protocols show high levels of entrainment, as compared to single 50 Hz synapse (bottom left) and synaptic bombardment (bottom right) protocols. Entrainment peaks occurred at every multiple of the intrinsic spiking frequency, here 10 Hz, that were not present for the two other conditions.

We further tested various configurations for gaussian noise stimulus and the two dynamic clamp protocols, since multiple parameters can be tuned to obtain the same spiking frequency but with potentially different variability in the ISI distribution. Therefore, we increased standard deviation of the gaussian noise centered at 0.552 nA, from half of its mean to 1, 2 and 4 times the mean. The resulting entrainment, quantified by PLV as mentioned previously, are depicted in Figure 3. Increasing standard deviation/noise power, globally decreased AEF entrainment, with fewer harmonics represented and even none for the highest level of noise.

**Figure 3.**
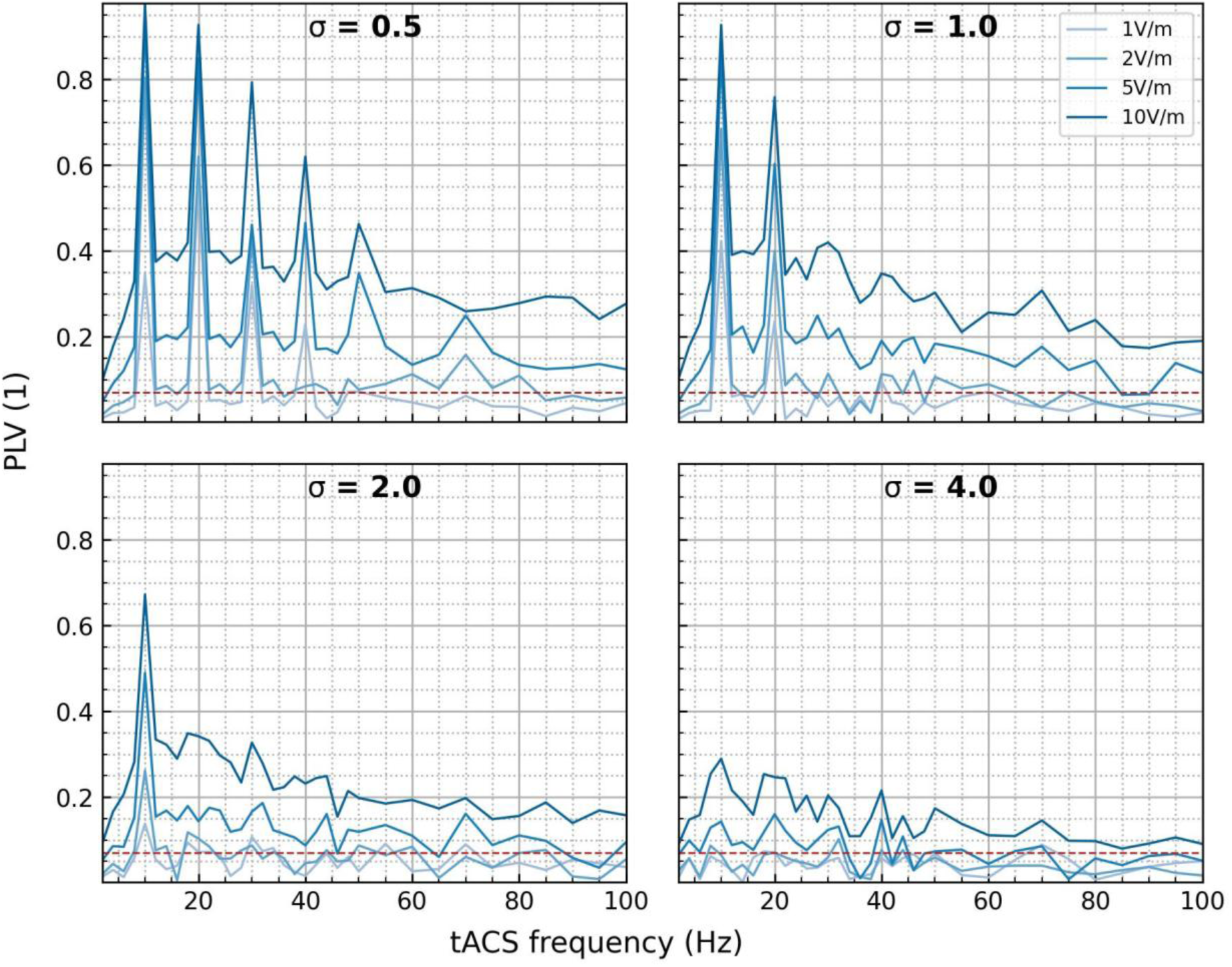
Layer 5 pyramidal cell’s response to gaussian noise stimulus with multiple standard deviation (STD) levels. PLV is depicted for a 0.5, 1, 2 and a 4 times mean STD, the mean being set to generate a 10 Hz cell’s activity. Increasing STD decreases Arnold tongue behavior, with a decrease in the number of peaks at harmonic of the firing rate.

Increasing the frequency of the single excitatory synaptic-like input of the first dynamic clamp protocol increased AEF sensitivity, with higher PLV over the entire spectrum (2-100 Hz) and the presence of a resonance at the spiking frequency of 10 Hz (Figure 4). This could be due to lower variations induced in the total current/conductance, since amplitudes of artificial excitatory postsynaptic potentials need to be decreased since its frequency of arrival increased, to keep spiking frequency at the same level. Interestingly, the AEF sensitivity followed the same trend as for gaussian noise protocols with the 4xmean standard deviation with higher values.

**Figure 4.**
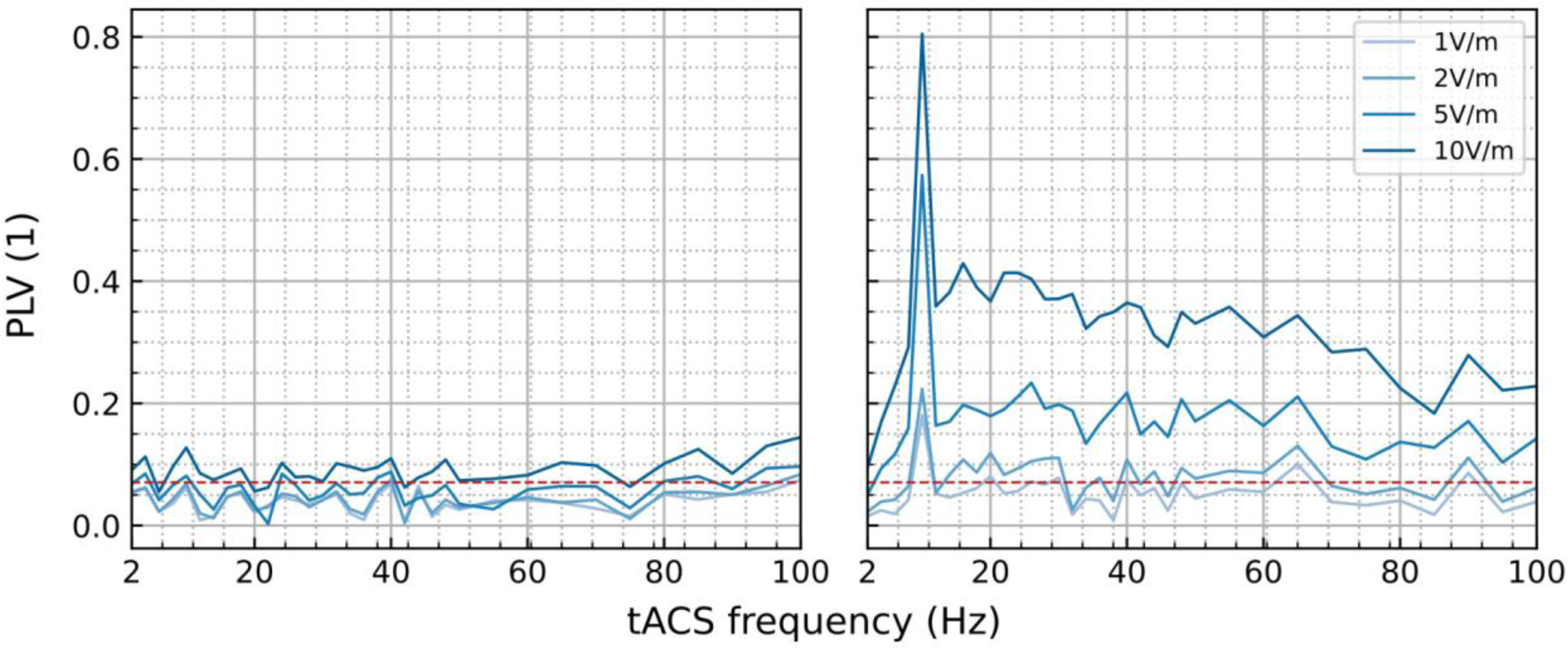
Impact of input synaptic frequency using one excitatory synapse stimulus at 50 Hz and 3000 Hz. PLV increased drastically at 3000 Hz, where a 5 V/m alternating electric field entrained activity in the [2–100] Hz range, with a clear peak at the firing rate (10 Hz). 50 Hz excitatory synaptic input entrainment did not show a clear pattern with a smaller entrainment, PLV being above significance threshold at only 10 V/m for most frequencies.

Regarding synaptic bombardment, tunable parameters were the mean and std of the gaussian process for both excitatory and inhibitory bombardments, since the time constant were kept as reported *in-vitro* measurements (Destexhe, 2001). Two different configurations were tested for the tested pyramidal cell: a 1:4 excitatory/inhibitory ratio (mean/std) and the 1:1 ratio that was also used in inhibitory models. Increasing the excitatory/inhibitory ratio by 4 increased the pyramidal cell’s sensitivity to AEF as illustrated in Figure 5, where PLV was above the significance threshold for all frequencies at the 10 V/m amplitude.

**Figure 5.**
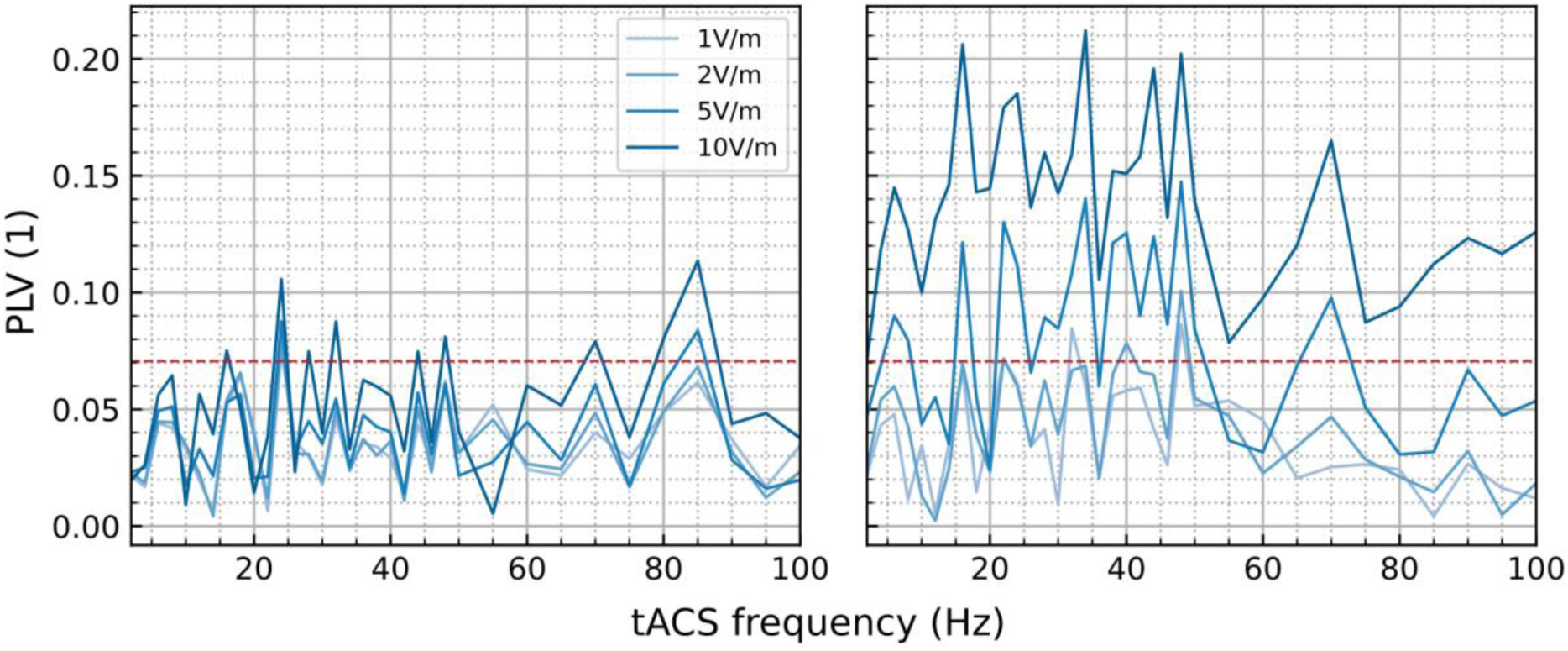
Impact of excitation/inhibition ratio in the synaptic bombardment protocol. The default ratio was 1:4 for the L5 PC (left) highlighted significant entrainment for several frequencies for a 10 V/m AES while the 1:1 ratio (right) showed an increase in cell’s sensitivity to the AEF with a significant entrainment at all frequencies for the same amplitude. No clear relationship was observed, with only a minor drop of entrainment when stimulation frequency is increased above 50 Hz for the 1:1 ratio, despite a peak at 70 Hz.

### Electric field sensitivity of inhibitory cells increases with frequency and is sensible to noise

The four tested GABAergic cells, one for SST and VIP types and two of the PV type, exhibited resonant sensitivity to AEF when using a monophasic current clamp (Figure 6). However, for all other protocols, inhibitory cells had the same frequency response, with an increased entrainment when ACS frequency increased, in line with previously reported results (Gaugain *et. al*, 2025). While PC exhibited the smallest ACS effect, GABAergic cells exhibited their highest among gaussian noise and single artificial synapse protocols. These results highlight a higher sensitivity to noise for GABAergic cells for current clamps, but also that dynamic clamps (artificial conductances) could reproduce frequency-dependent sensitivity to ACS observed when using reconstructed synapses.

**Figure 6.**
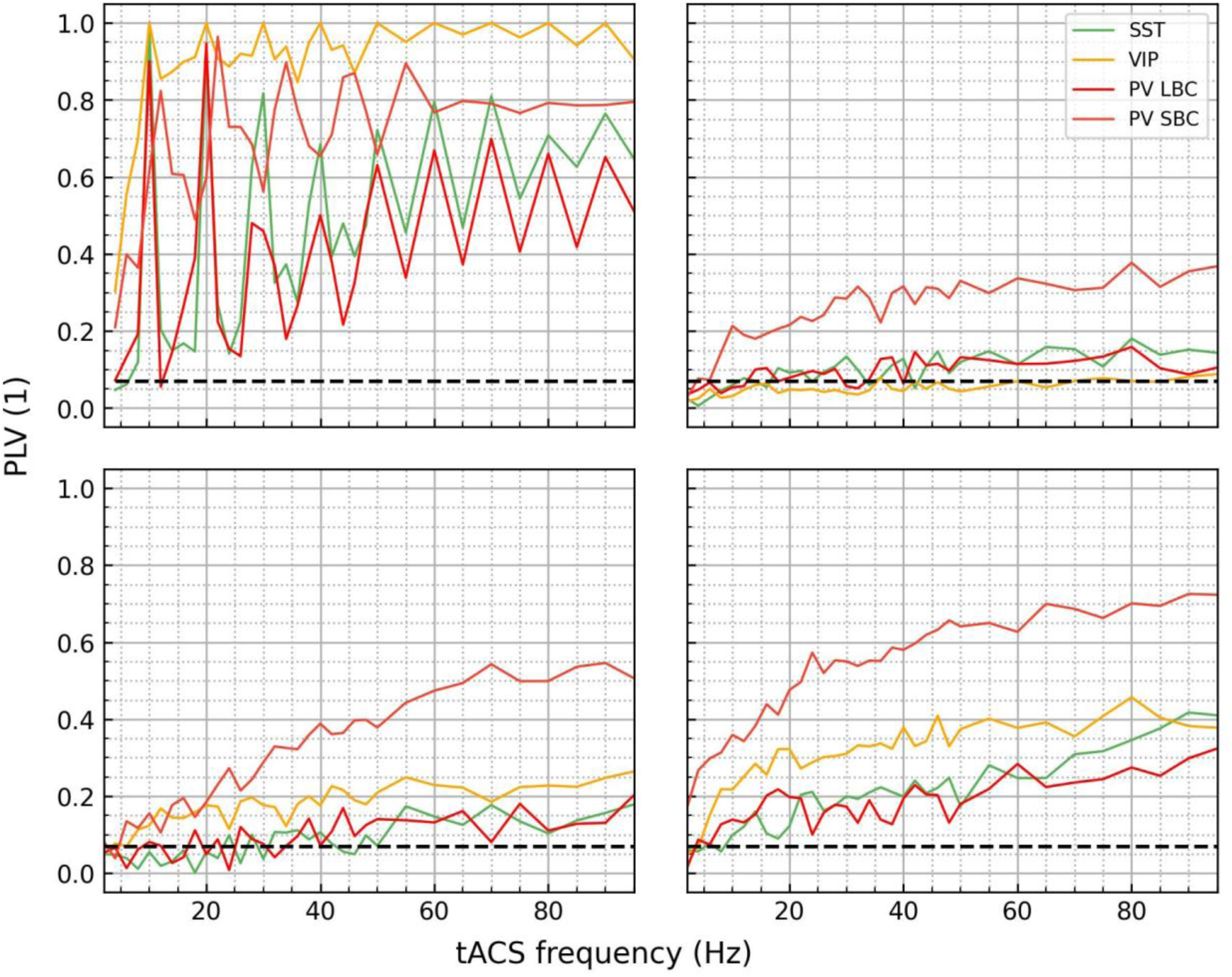
GABAergic cells’ response to the four tested clamp protocols during alternating electric field stimulation at 10 V/m for 2–100 Hz stimulation frequencies. Only monophasic current clamps (top left) produced high levels of entrainment with harmonics as compared to other protocols. In other clamp protocols, entrainment increased with stimulation frequency, with lower PLVs and increase for the gaussian noise current clamp protocol (top right) than for single synaptic current (bottom left) and the highest entrainment obtained with synaptic bombardment (bottom right).

## Discussion

To the best of our knowledge, this study is the first to investigate the input-dependent neuronal sensitivity to alternating current stimulation with precision, using a morphologically realistic modeling approach. These results are also transferable to all stimulation modalities mediated by electric fields, such as transcranial alternating magnetic field (Legros et al., 2024), even if the most commonly used technology is tACS. We showed that the measurement of such sensitivity to ACS is strongly impacted by statistics of the input used to drive neuronal activity, which has significant implications regarding the experimental investigation of tACS mechanisms. We showed that monophasic current clamps do not reproduce *in-vivo*-like sensitivity to ACS as observed in a previous study (Gaugain et al., 2025) and experimental *in vivo* conditions (Krause et al., 2019; Johnson et al., 2020), and produced Arnold-tongue-like behavior with resonances which is supposed to be a network behavior (Ali et al., 2013; Negahbani et al., 2018; W. A. Huang et al., 2021). Dynamic clamp protocols did not exhibit this unexpected behavior, and levels of sensitivity were closer from the previously observed ones, with GABAergic frequency-dependency closer to what was reported (Gaugain et al., 2025).

Single-cell response to an AEF depends on the cell type, as highlighted previously (W. A. Huang et al., 2021; Lee et al., 2024; Gaugain et al., 2025) in all input conditions, however levels of entrainments and frequency-dependency were input-dependent. This input dependency was previously suggested with supra-threshold paired pulse stimulation, showing increased effect of tACS on inhibitory neurons when pulse delivery frequency was increased (Khatoun et al., 2017). Interestingly, inhibitory neurons tended to have higher sensitivity to higher ACS frequencies when using gaussian noise inputs and synaptic-like inputs, consistently with using reconstructed multiple synapses (Gaugain et al., 2025). This higher frequency sensitivity was also present in monophasic currents, with higher harmonics present at 1 V/m (see figure S1), but this Arnold tongue was present only for a monophasic input. Altogether, these results strengthen further the tendency of inhibitory recruitment when using high frequency tACS. In contrast, layer 5 pyramidal cells, the most sensitive cells (Tran et al., 2022a; X. Huang et al., 2024; Gaugain et al., 2025) – due to their elongation along the neocortex – did not exhibit frequency preferences in either 50-Hz single synaptic process or synaptic bombardment. This result does not replicate the frequency preference found previously with reconstructed synapses. Neither did the monophasic current clamp, which exhibited clear resonances, with tremendous entrainment, far from the one reported in other studies (Tran 2022, Gaugain 2025) and experimental measurements (Krause et al., 2019; Johnson et al., 2020). Interestingly, this is also true for the 3000 Hz single synaptic process and the lowest gaussian noise intensity, where the pyramidal cell had only one resonance at the neuron firing frequency (10 Hz).

The reported input-dependent cell’s response to AEF may also impact network response and sensitivity, therefore their effect depends on the state of the network, which correlates with the findings of (Krause et al., 2022b) and a more recent report (Tran et al., 2025). Specifically, the L5 PC sensitivity, in the case of synaptic bombardment, depended on the excitation/inhibition balance, with a lower response with higher inhibitory synaptic activity. Here, only the statistic of the waveform was used but inhibitory shunting impact as well in *in vivo* condition (Paulus & Rothwell, 2016).

These findings highlight the need to take caution with protocol design when aiming to measure neurons’ sensitivity to AEF to avoid bias in results and conclusions about the involved underlying mechanisms. In this line, assessing this sensitivity to ACS might require natural synaptic inputs or stimulus based on the statistic of recorded natural activity/inputs, in an effort to reproduce it accurately during further single-cell measurements. Such an approach would improve our understanding of the reported *in vivo* results (Johnson 2019, Krause 2020, Krause 2022) that showed variability in results and one reporting endogenous activity-dependent sensitivity (Krause 2022). Finally, one limitation of using any somatic-clamp protocol is their inability to incorporate the contribution of dendritic processes, but this could serve as evidence and the quantification of the dendritic contribution to ACS effects, as well as synaptic plasticity.

## Acknowledgments

We acknowledge support of the SP-STIM project by the Labex COMINLABS (No. ANR-10-LABX-07-01).

## Supplementary

**Figure S1:**
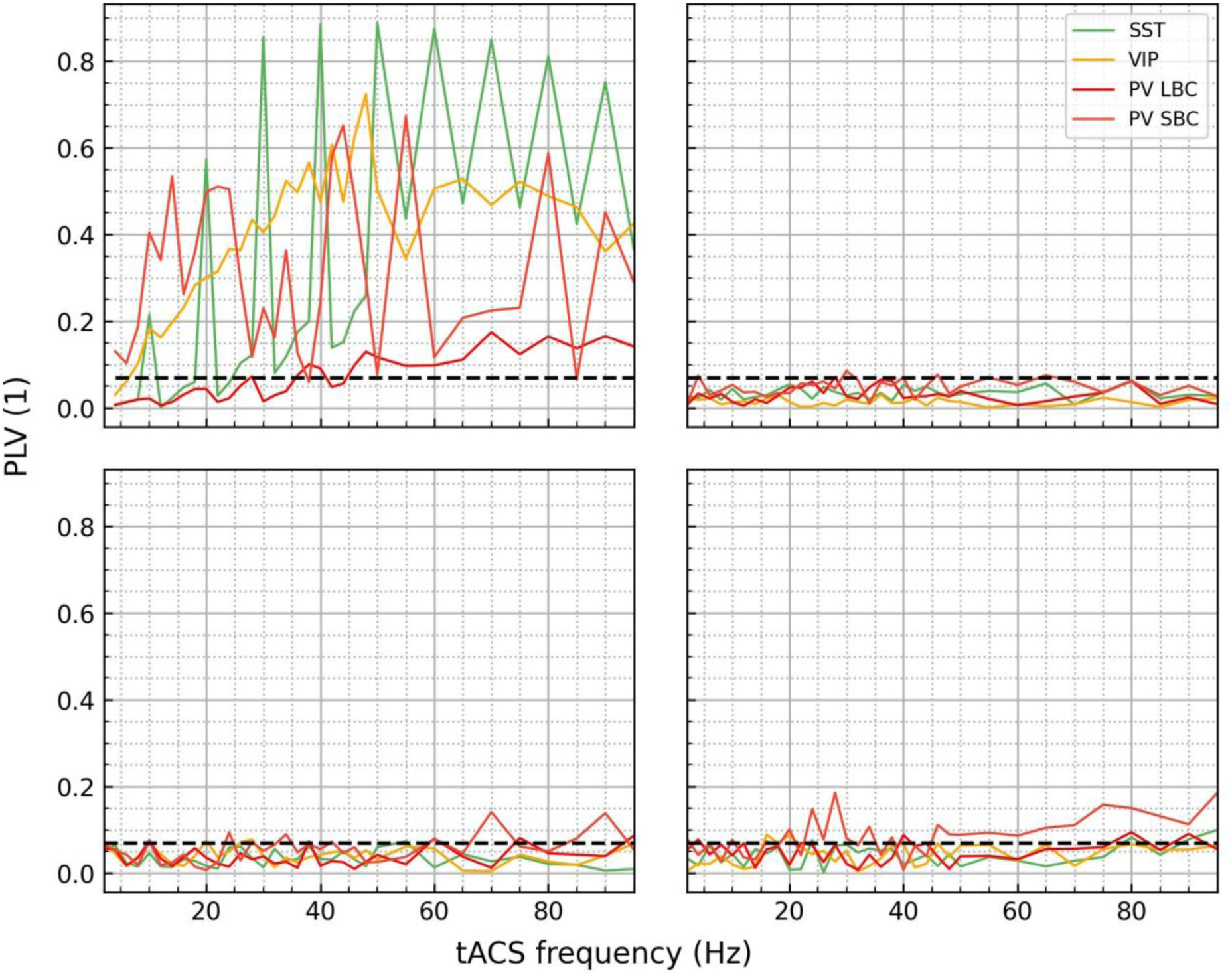
GABAergic cell’s entrainments for all the four protocols during exposure to an electric field magnitude of 1 V/m. Legend and formatting are similar to Figure 6.

**Table S1:** Parameters for each clamp stimulation. Icc is the current in nA delivered during the somatic monophasic current clamp protocol as well as the mean of the gaussian noise for the second one. *w*_50*Hz*_, *w*_3000*Hz*_ are the weight of the single synapse for triggering at 50 and 3000 Hz respectively for the single synapse protocol and γ 10 Hz is the mean of the gaussian noise for the synaptic bombardment protocol, considering half this value for the standard deviation (4 times the values for excitatory conductance for pyramidal cells).

| cells | lcc 10 Hz<br>(nA) | $w_{50Hz}$ 10 Hz<br>(1) | $w_{3000Hz}$ 10 Hz<br>(1) | $\gamma$ 10 Hz<br>(1) |
| --- | --- | --- | --- | --- |
| L5_TTPC2_cADpyr232_1 | 0.552 | 0.01236 | 2.174 e-04 | 0.0319 |
| L4_SBC_cACint209_1 | 0.029 | 0.000602 | 1.165 e-05 | 4.489 e-04 |
| L4_LBC_cACint209_1 | 0.04747 | 0.000091 | 1.856 e-05 | 5.96 e-04 |
| L4_MC_cACint209_1 | 0.04743 | 0.00108 | 1.965 e-05 | 5.438 e-04 |
| L4_BP_bNAC219_1 | 0.01812 | 0.000416 | 6.698 e-06 | 3.80 e-04 |

